# The Patent and Literature Antibody Database (PLAbDab): an evolving reference set of functionally diverse, literature-annotated antibody sequences and structures

**DOI:** 10.1101/2023.07.15.549143

**Authors:** Brennan Abanades, Tobias H. Olsen, Matthew I. J. Raybould, Broncio Aguilar-Sanjuan, Wing Ki Wong, Guy Georges, Alexander Bujotzek, Charlotte M. Deane

## Abstract

Antibodies are key proteins of the adaptive immune system, and there exists a large body of academic literature and patents dedicated to their study and concomitant conversion into therapeutics, diagnostics, or reagents. These documents often contain extensive functional characterisations of the sets of antibodies the describe. However, leveraging these heterogeneous reports, for example to offer insights into the properties of query antibodies of interest, is currently challenging as there is no central repository through which this wide corpus can be mined by sequence or structure.

Here, we present PLAbDab (the Patent and Literature Antibody Database), a self-updating repository containing over 150,000 paired antibody sequences and 3D structural models, of which over 65,000 are unique. Each entry in the database also contains the title and authors of its literature source. Here we describe the methods used to extract, filter, pair, and model the antibodies in PLAbDab, and showcase how PLAbDab can be searched by sequence, structure, or keyword.

PLAbDab uses include annotating query antibodies with potential antigen information from similar entries, analysing structural models of existing antibodies to identify modifications that could improve their properties, and compiling bespoke datasets of antibody sequences/structures known to bind to a specific antigen. PLAbDab is freely available via Github (https://github.com/oxpig/PLAbDab) and as a searchable webserver (https://opig.stats.ox.ac.uk/webapps/plabdab/).

## INTRODUCTION

Antibodies are by far the most successful type of biotherapeutic, with over 100 approved by the FDA and many more in the advanced stages of clinical development (Kaplon et al., 2023, Raybould et al., 2020). Their high specificity and affinity also make them a valuable tool in many areas of medical and scientific research. For example, antibodies are used routinely in diagnostic assays (Espejo et al., 2020), and to better understand the effects of vaccination on the immune system (Pollard and Bijker, 2021).

The variable region in antibodies (Fv) that is responsible for antigen binding, has a conserved global structure. It is formed by two immunoglobulin domains, the heavy (VH) and light (VL) chain variable domains. The binding site is divided between both chains, and is concentrated in six hypervariable loops, three on each chain, known as the complementarity-determining regions (CDRs). Among the CDR loops, the third CDR of the heavy chain (CDR-H3) is the most diverse in sequence and structure and often the primary contributor to antigen binding (Regep et al., 2017). However, the other five CDR loops and the relative orientation of the heavy and light chain have also been shown to affect binding (Gordon et al., 2023).

Next generation sequencing (NGS) has enabled researchers to take snapshots of the immune repertoire of an individual at a given point in time, leading to the generation of vast amounts of single chain antibody sequences. Efforts to compile this data has led to the creation of datasets such as Unpaired OAS (Kovaltsuk et al., 2018) and iReceptor (Corrie et al., 2018) which contain the sequence of the heavy or light variable domains for billions of antibodies. Paired VH-VL sequence data is more expensive to generate and currently just over a million paired antibody sequences can be found in Paired OAS (Olsen et al., 2021). However, with the binding site in antibodies sitting across both chains, paired data gives a more complete picture of how and to what an antibody binds (Jaffe et al., 2022, Shrock et al., 2023).

Although NGS data has proven invaluable to compare repertoires between individuals, it provides little information on the functions of individual sequences within a repertoire. However, there also exists a large number of smaller scale studies, each one dedicated to investigating a small number of antibodies. When combined, the antibodies from these studies amount to a large number of sequences with rich metadata. There are a number of databases that aim to compile subsets of this data, for example SAbDab (Dunbar et al., 2014, Schneider et al., 2022) for antibodies with resolved crystal structures, Thera-SAbDab for antibody therapeutics (Raybould et al., 2020), CoV-AbDab for COVID-19 binding antibodies (Raybould et al., 2021), or PAD for unpaired antibody sequences from patents (Krawczyk et al., 2021). Paired antibody sequences with information on their epitope can also be obtained from IEDB (Fleri et al., 2017).

Here we present PLAbDab, a database containing 150,000 paired antibody sequences from over 10,000 small scale studies. PLAbDab is larger than any other non-NGS database of paired antibody sequences by at least an order of magnitude. We make the data freely available and provide methods to rapidly search it by either sequence identity using KA-search (Olsen et al., 2022a), structural similarity (Robinson et al., 2021, Spoendlin et al., 2023), or by keywords in the title of the study. Each of the sequences comes with a direct link to its source material, making it easy to obtain additional information about any antibody of interest.

## METHODS

### Collecting unpaired antibody sequences

The majority of data in PLAbDab is extracted from the Protein database of NCBI (Sayers et al., 2021). The BioPython Entrez module (Cock et al., 2009) is used to query the database for entries containing the words “antibody”, “antibodies”, “immunoglobulin”, “scfv” or “bcr” in any of their fields. Entries with sequences longer than 1000 amino acids or shorter than 70 are removed at this stage. Entries containing the words nanobody or nanobodies are also removed. Around 2.5 million entries were returned from this search.

Sequences for each of these 2.5 million entries were then searched for antibody variable domain sequences using ANARCI (Dunbar and Deane, 2016). This resulted in around 530,000 potential antibody variable domain sequences from around 13,000 different sources.

### Creating paired antibody sequences from the unpaired data

For a large number of entries the metadata provides enough information to pair the heavy and light variable domains. For entries from the same literature source, VH-VL pairing was attempted using the following heuristics:

1. If there is only one VH and only one VL within a single entry, these were paired together.
2. If there is a unique non-common word that is found in the description of exactly one VH and one VL, these are paired together. Uncommon words are defined as words found in the description of less than 20 entries in the unpaired database.
3. Some entries contain information on the experimental source, such as the isolate or clone. If there is a unique VH and VL from the same experimental source they are paired together.
4. Entries from patents have sequence IDs by which they are referred to in the patent text. If the patent text mentions a VH and VL within the same paragraph, these are paired together. If a paragraph in the patent text mentions the same number of VH and VL entries, these are paired in the same order as they are mentioned.
5. If there is a single VH (or VL) from one source, this entry is paired with all other VL (or VH) entries from that source.
6. If there are the same number of VH and VL entries from one source, they are paired in order.

The described strategies for pairing VH-VL sequences have varying levels of accuracy. Twenty entries paired by each of the methods described above were randomly selected and manually checked to estimate the accuracy of each pairing method. Methods 1-4 achieved perfect accuracy on the twenty entry test set. Method 5 incorrectly paired one entry and Method 6 incorrectly paired two entries. Each paired entry is given a flag to indicate how it was paired, making it easy for users to filter for less accurate pairing methods. To further increase the coverage of the dataset, sequences from both SAbDab and Thera-SAbDab were also added to PLAbDab.

### Searching PLAbDab

To allow users to rapidly search the database for antibodies with a similar sequence, we implemented KA-Search (Olsen et al., 2022a). KA-search is able to carry out rapid and exhaustive sequence identity searches, allowing users to find similar sequences over the whole variable domain, the CDR loops, the CDR-H3 or a user defined region. The entire database can be queried with KA-Search in under 5 seconds on 5 CPUs.

Structurally similar antibodies can have similar functions, even if they are distantly related in sequence (Robinson et al., 2021, Spoendlin et al., 2023). To enable CDR structure-based searching, we model all paired sequences in the database using ABodyBuilder2 (Abanades et al., 2023). Entries missing more than eight residues at the start of the sequence were restored using AbLang (Olsen et al., 2022b) prior to modelling. Entries with the same sequence were only modelled once, in total 64,000 unique antibody models were generated.

To structurally search PLAbDab, query sequences are first modelled using ABodyBuilder2, without refinement. The refinement step was skipped as in order to boost speed with minimal compromise on backbone model quality. The framework of the predicted structure is then aligned to the structure of all other entries in PLAbDab with the same CDR loop lengths. Lastly, the carbon-alpha (C_*α*_) root-mean squared deviation (RMSD) over all CDR residues is computed and used to rank entries. Searching the database in this way takes around 10 seconds on 5 CPUs.

Finally, as each sequence in PLAbDab is linked to a patent or publication, we provide users the ability to search fields, such as the title of the study, using regular expressions.

## RESULTS

### Database statistics

Currently the total number of entries in PLAbDab is around 150,000, over 90% of which are paired with high confidence using methods 1-4, or comes from therapeutic or crystal structure entries. As can be seen in Figure 1A, the number of antibody sequences that could be collected by the PLAbDab methodology has been steadily growing since the early 2000s, with somewhere between 10,000 and 30,000 new antibody sequences being published each year for the last five years.

**Figure 1.**
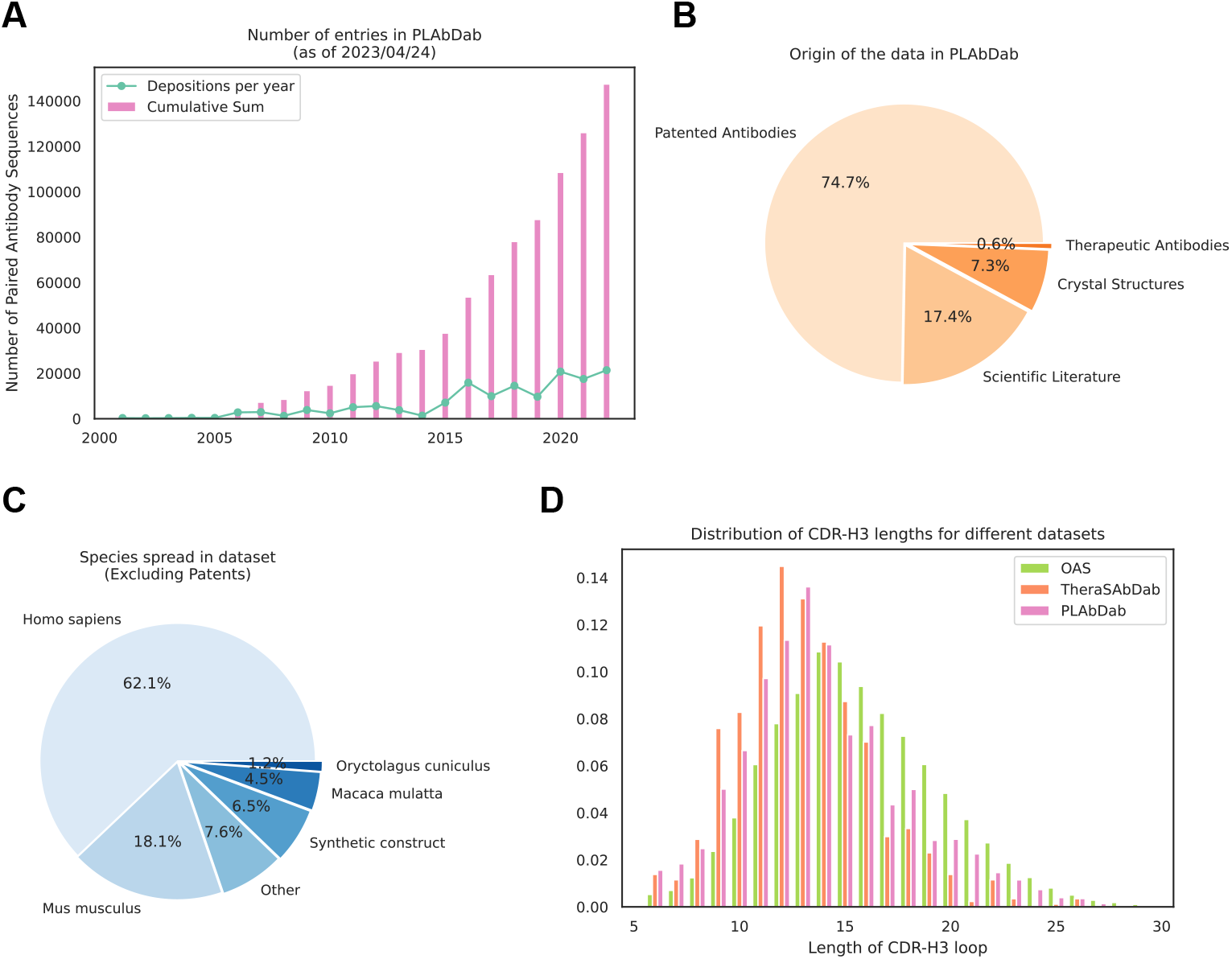
Summary statistics for PLAbDab. (A) Number of new antibody sequences each year and total number of antibody sequences that could be collected by the PLAbDab methodology. (B) The proportion of data in PLAbDab by type of source. (C) The proportion of antibodies from each species in non-patent entries. (D) Comparison of the distribution of CDR-H3 lengths in PLAbDab, Thera-SAbDab (Raybould et al., 2020), and OAS (Olsen et al., 2021).

Figure 1B shows the distribution of PLAbDab entries by source, around three quarters of entries come from antibody sequences described in patents, while less than 20% are derived from the scientific literature. This may be due to patents following a more standardised way of depositing antibody sequences. The rest of the entries in PLAbDab come either from solved structures in SAbDab (Dunbar et al., 2014),or from additional sequences in Thera-SAbDab (Raybould et al., 2020).

Patent applicants are not obligated to upload species information about their sequences to NCBI. This means that the majority of entries in PLAbDab lack a species annotation. However, species information does exist for the majority of the entries sourced from the academic literature. Across the sequences with species information, most entries are labelled as human, with a smaller number of sequences annotated as mouse, macaque or rabbit. (Figure 1C).

It has been observed that therapeutic antibodies tend to have shorter CDR-H3 loops than those seen in large repertoire studies, an observation that may be due to longer CDR loops leading to issues during therapeutic antibody development (Raybould et al., 2019, 2023). The average CDR-H3 loop length from antibodies in PLAbDab (around ∼14.0) falls somewhere between the average CDR-H3 loop length of OAS (around ∼15.6) and Thera-SAbDab (around ∼12.9) (Figure 1D).

### Searching PLAbDab

As described in the methods, PLAbDab allows users to search the database using a variety of methods. To demonstrate the sequence and structure search options, we compared the results of searching the database in four ways:

- For antibodies with a sequence identity of over 90% over the VH (VH identity).
- For antibodies with a sequence identity of over 90% for both the VH and VL (VH+VL identity).
- For antibodies with a C_*α*_ RMSD over the CDR loops of under 1.25Å to an ABodyBuilder2 model of the query (CDR structure).
- For antibodies with a C_*α*_ RMSD over the CDR loops of under 1.25Å to an ABodyBuilder2 model of the queryand sequence identity over the CDR loops of over 80% (CDR structure+identity).

We performed the above searches for three antibodies each of which target a different antigen. Antibody one binds programmed cell death protein 1 (PD-1), and was taken from the patent “Anti-PD-1 antibodies and methods of use thereof.”; antibody two binds Respiratory Syncytial Virus (RSV), sourced from the paper “Rapid profiling of RSV antibody repertoires from the memory B cells of naturally infected adult donors.” (Gilman et al., 2016); and antibody three is the therapeutic antibody Samalizumab, which binds the OX2 membrane glycoprotein (CD200). For each of the three queries we took the sequence and built an ABodyBuilder2 model for use in the structure searches. The result of using each search method described above to query PLAbDab with these antibodies is shown in Table 1.

**Table 1.** Results from searching PLAbDab with three different queries using four methods. For each method, the number of retrieved entries, the number of different sources the entries come from and the number of unique antibodies found is given. The value given in parenthesis is the number of entries or sources that are functionally consistent with the query. Details on the three query antibodies and the four searching methods are given in the main text.

| Search method | Retrieved (cons.) | Sources (cons.) | Unique (cons.) |
| --- | --- | --- | --- |
| PD-1 |  |  |  |
| VH identity | 576 (222) | 180 (41) | 258 (61) |
| VH+VL identity | 155 (132) | 39 (28) | 22 (16) |
| CDR structure | 227 (168) | 60 (26) | 46 (19) |
| CDR structure+identity | 127 (127) | 29 (29) | 14 (14) |
| RSV |  |  |  |
| VH identity | 2 (2) | 1 (1) | 2 (2) |
| VH+VL identity | 2 (2) | 1 (1) | 2 (2) |
| CDR structure | 29 (29) | 4 (4) | 23 (23) |
| CDR structure+identity | 4 (4) | 1 (1) | 4 (4) |
| CD200 |  |  |  |
| VH identity | 13 (13) | 4 (4) | 2 (2) |
| VH+VL identity | 13 (13) | 4 (4) | 2 (2) |
| CDR structure | 1175 (95) | 259 (15) | 440 (50) |
| CDR structure+identity | 37 (37) | 5 (5) | 5 (5) |

The PD-1 case study shows the benefits of having the paired heavy and light chain variable domain sequence. While a sequence identity search over the VH finds antibodies binding to the same epitope around 25% (61 out of 258) of the time, including the VL improves the accuracy to around 75% (16 out of 22). The fact that there are many highly identical heavy chain sequences that bind to different epitopes suggests that, for this antibody, the light chain may contribute significantly to binding. To validate this, we analysed the crystal structure of an antibody bound to RSV which is returned by all four search methods (PDB ID: 5GGR). According to Arpeggio (Jubb et al., 2017), the paratope is evenly split between the heavy and light chain, with eight heavy chain and seven light chain residues found to interact with the antigen.

Searching PLAbDab for entries similar to our PD-1 target by CDR structure returns antibodies that target the same epitope around 40% of the time (19 out of 46), but many of them are from different studies than those found by sequence search. All the entries returned by a CDR structure plus sequence identity search target the same epitope as our query.

For the RSV binding antibody, there are only two entries in PLAbDab with greater than 90% sequence identity over the VH. Searching PLAbDab for entries with a similar CDR structure returns 23 unique antibodies which bind the same epitope. This suggests that this antibody may require a very specific CDR loop conformation to bind the RSV Fusion Glycoprotein (Mukhamedova et al., 2021) at that specific site. Two of the retrieved entries belong to antibodies with resolved crystal structures (PDB IDs: 7LUC and 7LUD), one of which is bound to the antigen. Figure 2 shows the similarity of these structures to the ABodyBuilder2 predicted structure of the RSV binding antibody query.

**Figure 2.**
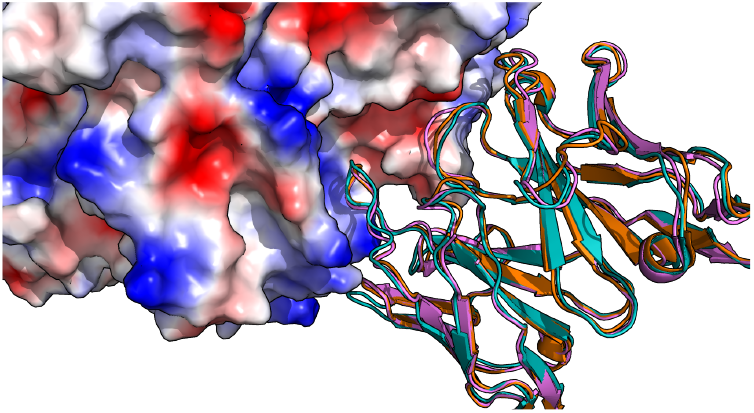
Crystal structures of anti-RSV antibodies aligned to structural model of query. A cartoon representation of the ABodyBuilder2 model for the query antibody is shown in purple with the crystal structure for 7LUC shown in blue and the crystal structure for 7LUD shown in orange. The surface of the RSV antigen is coloured based on electrostatics using PyMol (DeLano et al., 2002).

Only two unique antibody sequences in PLAbDab have a VH sequence identity of over 90% to Samalizumab. A structure search using Samalizumab as a query results in over a thousand antibodies that bind a very diverse set of epitopes. In Figure 3 we show a selection of antibodies with very similar CDR backbone structures to Samalizumab binding to four very different epitopes on different antigens. This indicates that the backbone structure of the CDR loops is likely not the primary driver of binding specificity for this antibody. However, when adding an 80% sequence identity cutoff over the CDR loops to the structure search, we are left with five unique antibody sequences all of which bind the same epitope.

**Figure 3.**
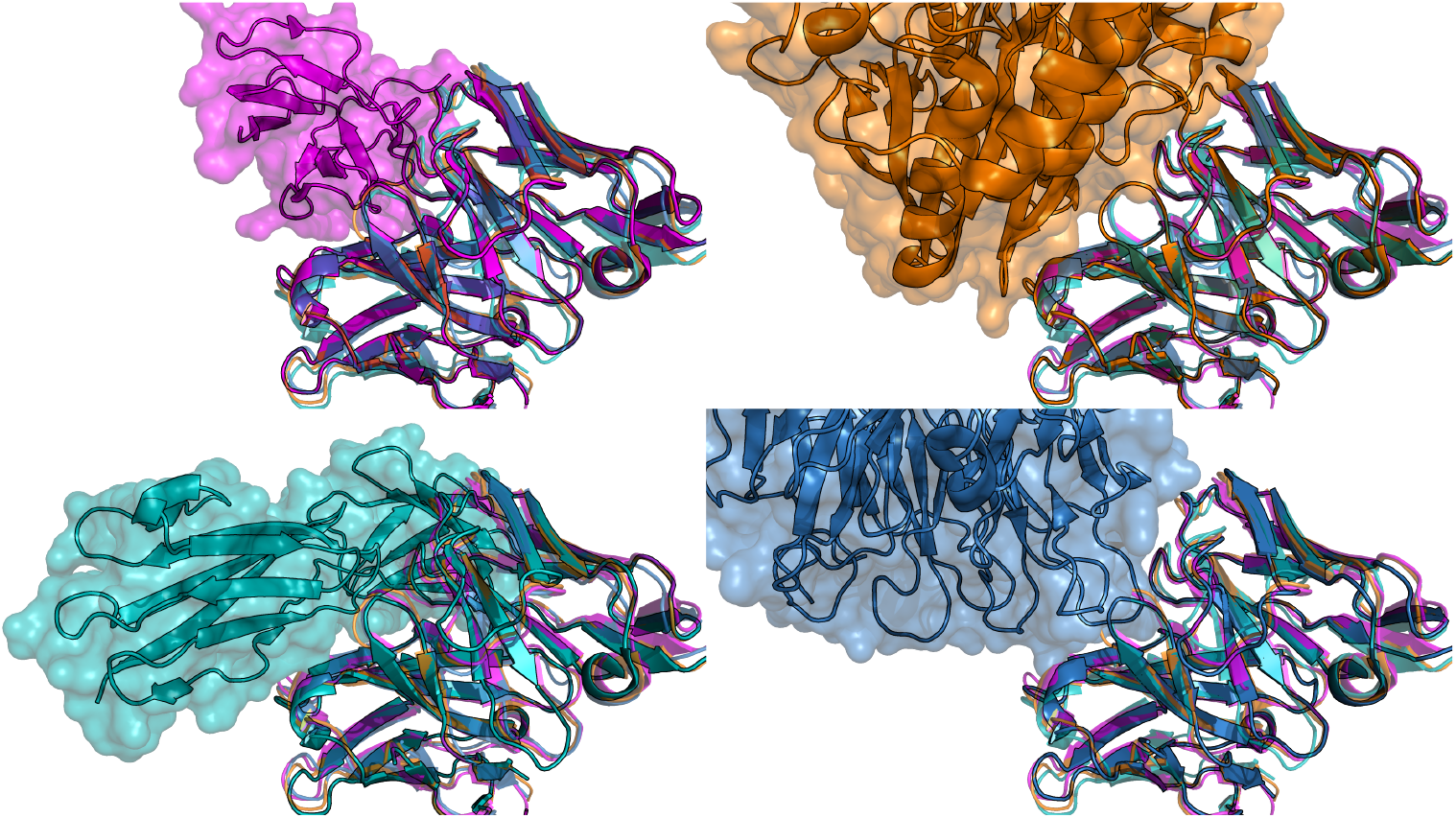
Crystal structure of four antibodies with very similar structure that bind very different antigens. The crystal structure of all four aligned antibodies is shown in each figure (4AL8 in purple, 7XY8 in green, 7LA4 in blue, and 7BBJ in orange). On the top left they are shown in complex with the antigen of 4AL8 (Dengue virus DII protein); top right 7BBJ (CD73); bottom left 7XY8 (Emmprin CD147); and bottom right 7LA4 (Integrin *α*_IIb_*β*_3_).

These five case studies highlight the utility of PLAbDab’s data and various search methods to identify antibodies with similar function to a query. The case studies also highlight that any computational search should be further investigated via both the meta data in PLAbDab and, if possible, experimental means.

### Using PLAbDab to Generate New Datasets

As PLAbDab contains a large number of antibody sequences targeting a diverse set of antigens, it can be used to generate antigen-specific antibody libraries. For example, searching PLAbDab for entries containing the keywords “ebov” or “ebola” in their title returns almost 1,500 unique antibody sequences from 56 sources. Using the keywords “hiv” or “immunodeficiency” finds over 6,200 entries with over 3,800 unique antibody sequences from over five hundred sources, while “autoimmune” and “autoantibody” returns over 400 unique antibody sequences from over 90 sources. Not every antibody sequence retrieved from a patent or paper will necessarily bind the searched target, but such prefiltered sets provide a very valuable starting point towards generating an antigen specific library to a target of interest.

To benchmark how effective searching the titles of the sources of PLAbDab entries by keywords is at retrieving relevant antibodies, we manually inspected a random set of 100 antibodies from different studies and found that 88 are true HIV binders. Of the non-HIV binders, four were anti-idiotype antibodies, three bound CD4, three bound other antigens and two were incorrectly paired.

## DISCUSSION

We present PLAbDab, a large database of paired antibody sequences from patents and papers. Each entry in the database contains a link to its original source material, which relates the paired sequences to useful functional information. PLAbDab can be automatically updated without any manual input, making it straightforward to keep up to date with the latest publications and patents.

To demonstrate the utility of PLAbDab, we explore different search methods for identifying antibodies with similar binding properties to a given antibody of interest. Our results demonstrate that for different antibodies, different types of search yield the best results. In all three cases investigated, CDR structure alongside CDR sequence identity gave the most accurate results, but in most cases, this also missed functionally cognate antibodies. The inclusion of the light chain in any search always improved accuracy, in agreement with recent studies that have highlighted the importance of the light chain residue motifs for antigen binding (Shrock et al., 2023), and restricted heavy and light chain gene pairs observed among functional antibodies (Jaffe et al., 2022). In one case searching by structure provides a link that is not immediately apparent from sequence search alone. This finding is consistent with recent work which uses structural information for epitope binning (Robinson et al., 2021, Spoendlin et al., 2023).

One of the main challenges in developing PLAbDab, and the area with most potential for improvement, is the pairing of heavy and light chains based on metadata. Although the strategy described in this paper accurately pairs a large number of antibody sequences, there are still many for which pairing was not possible. The PLAbDab generation code also relies on authors publishing their antibodies to databases such as NCBI (Sayers et al., 2021) or the PDB (Berman et al., 2000), and although many authors do submit their sequences to these databases, there remains a large body of antibody sequence data shared in the text, figures, or tables of papers.

Nevertheless, PLAbDab already contains over 60k unique annotated antibody sequences providing an invaluable resource to the antibody research community. PLAbDab can also be used to generate antigen specific libraries that could be used to train novel machine learning models or as a starting point for the development of novel therapeutics. We have made PLAbDab freely available to download and query via a webserver (https://opig.stats.ox.ac.uk/webapps/plabdab/) and the code is available to download via github (https://github.com/oxpig/PLAbDab).

